# Annexin A1 tripeptide mimetic increases sirtuin-3 to augment mitochondrial function and limit ischemic kidney injury

**DOI:** 10.1101/2020.12.11.421859

**Authors:** Hagir Suliman, Qing Ma, Zhiquan Zhang, Jiafa Ren, Benjamin T. Morris, Steven D. Crowley, Luis Ulloa, Jamie R. Privratsky

**Author notes:** **Correspondence to:** Jamie R. Privratsky, M.D., Ph.D. Department of Anesthesiology Duke University Medical Center DUMC Box 3094, Durham, NC 27710, E. **Declaration of interest:** ZZ and QM are coinventors on patents filed through Duke University on the therapeutic use of Annexin A1 tripeptide (ANXA1sp).

## Abstract

**Background:** Acute kidney injury (AKI) is one of the most common organ failures following surgery. We have developed a tripeptide mimetic (ANXA1sp) of the parent annexin A1 molecule that shows promise as an organ protectant limiting cellular stress; however, its potential as a kidney protective agent remains unexplored, and its mechanism of action is poorly understood. Our hypothesis was that ANXA1sp would limit kidney injury and improve mitochondrial function following surgical ischemic kidney injury.

**Methods:** In blinded fashion, wildtype mice were assigned to receive vehicle control or experimental drug (ANXA1sp) 1 hour prior to and 1 hour after kidney vascular clamping. Our primary outcome was assessment of kidney injury and function by measurement of serum creatinine and blood urea nitrogen (BUN) and histologic injury scoring of kidney tissue sections. Immunofluorescence microscopy, real-time PCR and western blot were used to assess cell death, oxidative stress, and mitochondrial biomarkers. An in vitro model of oxygen-glucose deprivation in immortalized kidney tubule cells was used.

**Results:** ANXA1sp given prior to and after ischemic kidney injury abrogated ischemic AKI. ANXA1sp further limited kidney cell death and oxidative stress following ischemia. ANXA1sp significantly improved markers associated with mitochondrial DNA repair and mitochondrial biogenesis. ANXA1sp upregulated expression of the mitochondrial protectant sirtuin-3 (SIRT3) in the mitochondria of kidney tubular cells. Silencing of SIRT3 limited ANXA1sp-mediated protection against hypoxic cell death.

**Conclusions:** ANXA1sp limits kidney injury through upregulation of SIRT3 and consequent preservation of mitochondrial function. ANXA1sp holds considerable promise as a perioperative kidney protectant prior to ischemia inducing surgery and/or kidney transplantation.

## Introduction

Acute kidney injury (AKI) is one of the most common forms of organ injury occurring in up to 5% of all hospitalized patients, 10-30% of post-surgical patients (49), and 30% of critically ill patients (42). AKI increases morbidity and mortality and results in longer ICU and hospital stays, leading to increased hospital costs (15). Even small changes in serum creatinine are associated with increased perioperative and long-term mortality (17, 19, 20). Despite its significant morbidity and mortality, there are currently no therapeutic modalities to prevent or treat AKI once it occurs. Thus, novel therapeutic modalities are needed.

Due to its high metabolic demands and oxygen consumption, the kidney is particularly susceptible to metabolic and oxidative stress (2). Kidney tubule cells are rich in mitochondria that are required for efficient ATP production via oxidative phosphorylation (2). Kidney tubular cells depend primarily on mitochondrial energy production making them sensitive to mitochondrial dysfunction, and mitochondrial impairment in the kidneys can severely affect kidney health (2, 12). Several studies have suggested that mitochondrial damage and dysfunction contribute significantly to AKI development and impede kidney repair and regeneration (43, 44). For example, mitochondrial fragmentation, swelling, and inner cristae loss were observed in experimental models of ischemic AKI (29, 47, 48) even prior to overt kidney cell apoptosis (5). Mitochondria are central to the regulation of both cellular metabolism and the integration of pathways that lead to cell death within the kidney. As such, targeting mitochondrial quality control is a promising therapeutic target. In this regard, sirtuin 3 (SIRT3) is a mitochondrial NAD^+^ dependent deacetylase that maintains mitochondrial integrity under conditions of cellular stress (16, 24, 30). In addition, SIRT3 has been shown to protect against toxic (26) and septic (54) AKI. Developing therapeutic agents that upregulate SIRT3 and protect the mitochondria, could have broad implications for kidney protection prior to AKI-inducing stimuli (i.e. surgery, transplantation) and during recovery following AKI.

Annexin A1 is a 37kD endogenous protein that is expressed mainly by immune cells and epithelial cells (21). Annexin A1 is a well-established pro-resolving, anti-inflammatory mediator (10, 21). As a result, peptide fragments of this molecule have been generated and shown to have protective anti-inflammatory properties in many disease states (38), including in kidney ischemia/reperfusion injury in rats (8). Our group has developed a specific small tripeptide fragment of the human annexin A1 molecule (ANXA1sp) that exerts potent biologic properties (52). Based upon the promising protective role of SIRT3 in toxic and inflammatory AKI (26, 54), we hypothesized that ANXA1sp would protect against ischemic AKI through upregulation of SIRT3, mitochondrial protection, and amelioration of tubular cell death. Here, we analyze how ANXA1sp treatment affects kidney injury, mitochondrial function, SIRT3 levels, and cell death pathways following ischemic AKI. These studies have important implications for kidney protection during surgery and prior to kidney transplantation.

## Materials and Methods

### Chemicals and Reagents

Annexin A1 tripeptide fragment (Ac-Gln-Ala-Trp) (ANXA1sp) was synthesized by GenScript Biotech (Piscataway, NJ) as previously described (52) and was reconstituted in DMSO and placed in individual doses. Ketamine, xylazine, and buprenorphine were purchased from Henry Schein animal health (Dublin, OH).

### Animal Experiments

All of the animal studies were approved by the Durham Veterans Affairs Medical Center (VAMC) Institutional Animal Care and Use Committee, performed at the Durham VAMC, and conducted in accordance with the National Institutes of Health Guide for the Care and Use of Laboratory Animals. Briefly, 129/SvEv 10-16-week-old male mice were obtained from Taconic Biosciences (Rensselaer, NY). Mice were fed standard chow diet.

#### Administration of ANXA1sp

Mice were randomly assigned to receive vehicle control or blinded experimental drug in every other fashion. The investigators performing surgery, experiments, and analyzing the data were blinded to the treatment groups until measurement of primary outcome for each experiment. Both DMSO Vehicle and ANXA1sp doses were reconstituted in saline and at 1 hour prior to clamp placement, 1mg/kg was given intraperitoneally (IP). The same treatment was given at 1-hour post-clamp removal.

#### *Ischemia/reperfusion* (I/R)

Mice were anesthetized with ketamine/xylazine. Mice were placed on a warming pad (Hallowell EMC, Pittsfield, MA) heated to 38°C by a Gaymar TP650 water pump. After aseptic prep, a midline dorsal incision was created, and blunt dissection was performed toward right kidney. The flank muscle and fascia above the right kidney was incised and the right kidney was exteriorized after which the renal pedicle was ligated with suture and the right kidney removed. After closure of fascia and muscle over right kidney, blunt dissection was performed toward left kidney. The flank muscle and fascia above the left kidney was incised and the left kidney was exteriorized. Adipose and connective tissue were carefully removed near renal vessels and a 800g pressure clamp (Fine Science Tools) was placed on the left renal pedicle. At end of ischemic time, the clamp was removed and reperfusion was confirmed by color change in kidney. The fascia and muscle layer and skin were closed by suture. The animal was given buprenorphine (0.1mcg/gm) in normal saline subcutaneously. The animal was then placed in cage with warming pad during recovery.

### Blood & Serum Analyses

Blood was collected from a cardiac puncture at the indicated time points, allowed to clot for 30 mins at room temperature, and centrifuged at 3,000 g for 10 minutes at 4°C. The primary outcome of our study was reduction in serum creatinine and BUN. Serum creatinine levels were measured by the University of North Carolina Animal Histopathology and Lab Medicine Core. Serum BUN was measured by kit (ThermoFisher) according to kit instructions.

### Histologic Analyses

Kidney tissues were removed and a cross-sectional segment obtained. The kidney segment was fixed with 10% neutral-buffered formalin (VWR 16004-128); embedded with paraffin; sectioned in 5mm sections by the Duke Research Immunohistology Laboratory.

#### Injury scoring

Sections were stained with PAS staining and scored by an experienced animal pathologist masked to experimental groups. Sections were graded according to a previously established scoring system (50): the percentages of tubules with cell lysis, loss of brush, border, and cast formation were scored on a scale from 0-4 (0, no damage; 1, 25%; 2, 25-50%; 3, 50-75%; 4, >75%). Histologic scores for each kidney were obtained by adding individual component scores.

#### 8-OHdG and citrate synthase immunofluorescence staining

Paraffin-embedded kidney sections (4 μm) were processed for immunostaining by deparaffinization in xylene and rehydration through a descending series of alcohols, washed extensively in 0.1 M PBS, blocked with 10% normal goat serum, and incubated in primary antibodies to 8-hydroxy-2’deoxyguanosine (8-OHdG) or citrate synthase diluted in 10% normal goat serum overnight at 4°C. followed by fluorescent-labeled secondary antibodies labeled with Alexa Fluor 488 green or Alexa Fluor 556 red. Nuclei were counterstained with DAPI (blue). Images were acquired on a Nikon E400 fluorescence microscope.

#### SIRT3 and mitochondrial complex IV immunofluorescence staining

After deparaffinization, kidney tissue sections were treated with 10 mM citrate buffer (pH 6.0) for antigen retrieval. After blocking with 10% normal goat serum and 0.1% BSA at RT for 1 hour, the sections were incubated with rabbit anti-SIRT3 (1:300; Cell Signaling Technologies, Danvers, MA) and mouse anti-COXIV (1:500; Santa Cruz Biotechnology, Santa Cruz, CA, United States) at 4°C overnight. The sections were then incubated with Alexa Fluor 488-conjugated goat anti-rabbit IgG (1:500; Invitrogen, Carlsbad, CA, United States) and Alexa Fluor 550-conjugated goat anti-mouse IgG (1:500; Invitrogen, Carlsbad, CA, United States) at RT for 1 hour. After washing with PBS, slides were prepared and mounted using UltraCruz™ Mounting Medium with DAPI (Santa Cruz Biotechnology, Santa Cruz, CA, United States) to detect nuclei. Images were captured on a Leica fluorescent microscope (Leica DM IRB, Germany) using a 20X/0.4 PH objective at 1.5-fold magnification, and the images were analyzed by NIH ImageJ software (version 1.51).

### RT-PCR

mRNA was isolated with an RNeasy Mini Kit (Qiagen, Germantown, MD) per kit instructions. A cross-sectional piece of kidney containing cortex and medulla was homogenized in Buffer RLT with 0.01% β-ME and further homogenized with Qiashredder columns (Qiagen 79654). mRNA concentration was measured by Nanodrop (ThermoFisher). The High Capacity cDNA Reverse Transcription Kit (Applied Biosystems 4368814) was used to synthesize cDNA according to manufacturer’s instructions. Gene expression levels of SIRT3 was determined by RT-PCR on an ABI7900HT machine (Applied Biosystems) using TaqMan primers (SIRT3, Mm00452131_m1, #4331182--ThermoFisher).

### Western Blot

A piece from flash frozen kidneys were homogenized in RIPA buffer containing cocktail protease and phosphatase inhibitors. Kidneys representative of mean serum creatinine for each group were used for Western blot samples. Total protein content was measured by the BCA assay. 20 μg total protein was loaded into SDS-PAGE gels and immunoblots were performed as previously described (39). Antibodies used include peroxisome proliferator-activated receptor-γ coactivator-1α (PGC-1α; 1:1,000; Cat. No. ab54481; Abcam); mitochondrial transcription factor A (Tfam; 1:1,000; developed in our laboratory); Pink1 (1:500; cat. no. ab23707; Abcam); PARK2 (1:500; Cat. No. sc-32282; Santa Cruz Biotechnology); light chain 3B protein (LC3I/II) tubulin (1:1,000; Cat. No. T-5168; Sigma-Aldrich), or mitochondrial porin (1:500; Cat. No. sc-8829), Drp1 (Santa Cruz Biotechnology, Dallas, TX), sirtuin 3 (#5490S, Cell Signaling Technologies, Danvers, MA).

### Cell culture

#### HK-2 cell culture

The human immortalized proximal tubule epithelial cell line HK-2 (ATCC, CRL-2190) was kindly provided by Dr. Tomokazu Souma and cultured in DMEM/F12 supplemented with 10%FBS, 1% penicillin/streptomycin, 1% Insulin-Transferrin-Selenium solution (Gibco).

#### Hypoxia exposure

Cells were exposed to ANXA1sp (10, 20 μM) for 1 hr. Cells were then subjected to 12 or 16 hr oxygen–glucose deprivation (OGD: DMEM no glucose, 92% N2/ 3% H2/ 5% CO2) in an anaerobic chamber (Coy Laboratories). Cells were lifted and analyzed for apoptosis/cell death per below.

Normoxia cells were treated with DMEM + F12 with no serum in a 37°C growth incubator with 95% air/5% CO2. Vehicle cells were treated with equal volumes of DMSO in the medium. Cells were harvested for further analysis.

### Cell Death Analysis

#### TUNEL staining

Apoptosis was determined by terminal deoxynucleotidyl nick-end labeling (TUNEL) per assay manufacturer’s protocol (Roche Diagnostics, Indianapolis, IN, United States). Paraffin-embedded sections (5 μm thick) were deparaffinized using xylene and descending grades of ethanol and incubated in a proteinase K working solution for 15-30 minutes at room temperature. Sections were then incubated with terminal deoxynucleotidyl transferase (TdT) for 1h at 37◦C and then rinsed with PBS. Slides were counterstained with DAPI and coverslipped using Dako Fluorescence Mounting Media (Agilent Technologies, Santa Clara, CA). Slides from three representative mice of each group were chosen based on nearness to mean of creatinine for each group. For each animal, five areas of the corticomedullary junction were imaged at 20X; images were post-processed within Image J with intensity parameters equal amongst all images; and nuclei were scored as positive if there was DAPI and TUNEL staining positivity in the same location. TUNEL positivity for each animal was reported as the number of TUNEL positive nuclei per 20X field averaged over the five images.

#### Cell death assay

HK-2 were treated with ANXA1sp and hypoxia per above. At end of experiment, the supernatant was removed to collect dead cells, cells lifted with TrypLE (Gibco), TrypLE inactivated with 10% FBS, and cellular fractions combined with supernatants. Cells were spun down (350g for 5 min), and pellet was resuspended in 100 ul media. Apoptosis was determined by Muse Annexin V & Dead Cell Kit (Luminex) per kit instructions.

### Statistical Analyses

Tests were performed with GraphPad Prism Software® (GraphPad Software, La Jolla, CA). The figures are representative of experiments that were repeated at least twice on different days, and the data are expressed as mean ±standard error of the mean (SEM). An unpaired Student’s t-test with Welch’s correction was used to compare two experimental groups. Comparisons with more than 2 groups were analyzed by one-way or two-way ANOVA as indicated in each figure legend with Sidak’s multiple comparisons test for post-hoc analysis to compare groups. Control group for animal experiments was Vehicle-treated, Sham mice. We did not perform *a priori* power analysis to determine sample size because we did not know the expected effect size of ANXA1sp on ischemic AKI. However, we attempted to perform each surgical experiment at least twice to target 12 mice/group, which is within range for a typical sample size for AKI in our laboratory (33, 51).

## Results

### ANXA1sp tripeptide prevents kidney injury following ischemia

Annexin A1 is a pro-resolving mediator (10, 21), and peptide fragments of annexin A1 have been generated and shown to have protective properties in many disease states (38). Here, we use a synthetic tripeptide of the N-terminal domain of the annexin A1 molecule (ANXA1sp) that retains the most potent anti-inflammatory and pro-resolving properties (52) in the setting of ischemic kidney injury to determine its effect on kidney protection. We hypothesized that pre-treatment of mice with ANXA1sp would dampen severity of AKI following ischemia. We injected mice with either vehicle or ANXA1sp 1 hour prior to ischemia, induced severe ischemia, and then injected mice with the same treatment 1 hour after reperfusion. In this ischemia model, we noted significantly less tubular injury (Figure 1A-C) and lower serum levels of creatinine (Figure 1D) and blood urea nitrogen (BUN) (Figure 1E) at 24 hours in ANXA1sp-treated mice. We concluded that ANXA1sp treatment ameliorated ischemic kidney injury.

**Figure 1.**
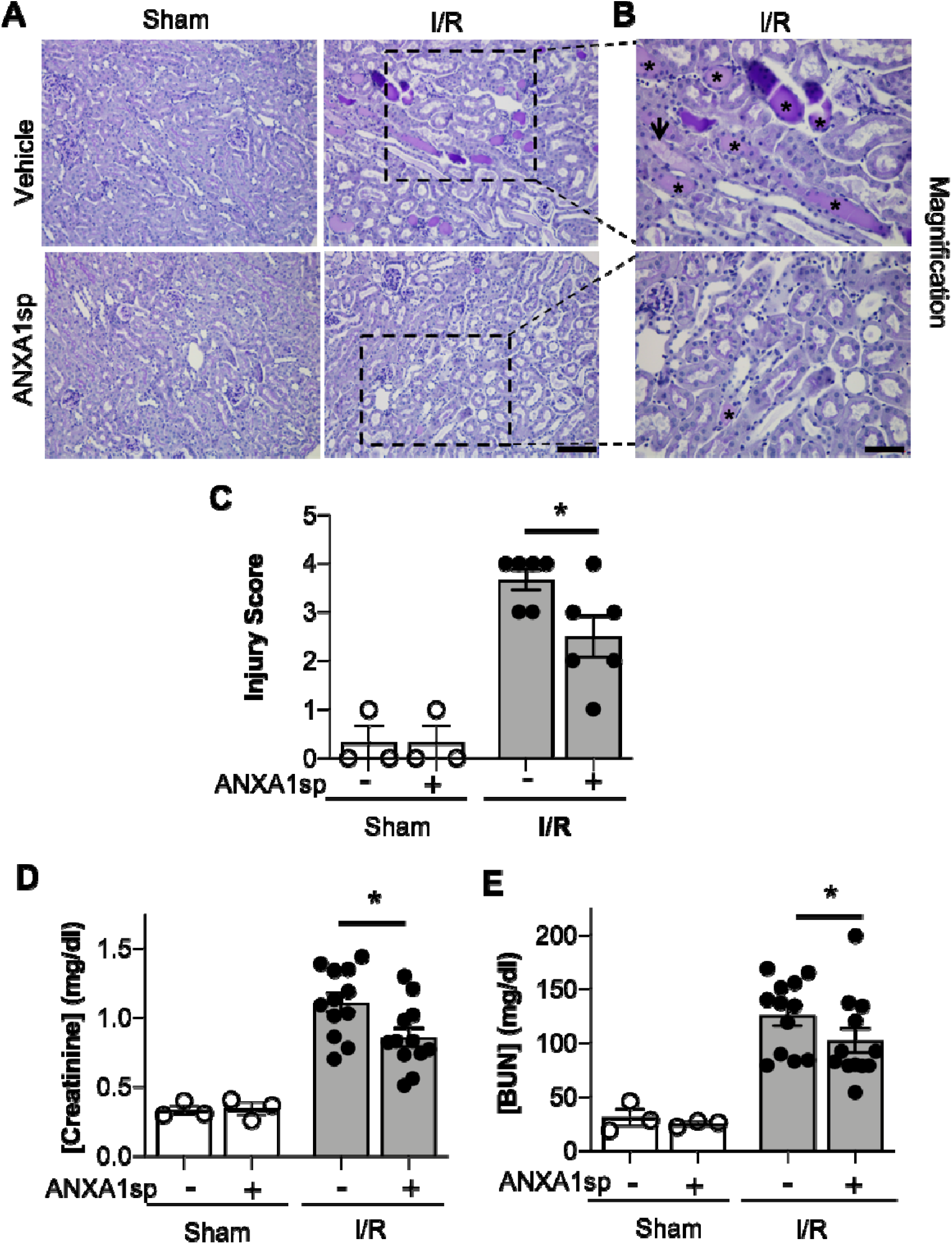
AnnexinA1 tripeptide (ANXA1sp) attenuates kidney injury following ischemia/reperfusion-induced kidney injury. (A) Mice were treated with either Vehicle or ANXA1sp and subjected to 33 minutes of unilateral ischemia and contralateral nephrectomy and then re-injected with Vehicle or ANXA1sp 1 hour after reperfusion. Representative periodic-acid Schiff (PAS)-stained kidney sections demonstrating increased injury in Vehicle, I/R group at Day 1 after ischemia (scale bar = 100 um). (B) Increased magnification of boxes in (A) to demonstrate increased histologic evidence of injury in Vehicle, I/R mice (asterisk: protein casts; arrowhead: tubule vacuolization) compared to ANXA1sp-treated mice (scale bar = 50 um). (C) Histologic injury scoring from (A) by observer blinded to experimental grouping (n=3 for Sham groups, n=6 for I/R groups). Six representative samples were selected based on mean creatinine of complete sample set for each group. ANXA1sp-treated mice display attenuated kidney injury. Statistical significance determined by two-way ANOVA (*p<0.05). (D, E) Serum creatinine (D) and blood urea nitrogen (BUN) (E) were measured. ANXA1sp-treated mice display ameliorated AKI compared to Vehicle-treated mice (n=3 for Sham groups, n=12 for I/R groups). Graphs display mean +/− SEM. Statistical significance determined by two-way ANOVA (*p=0.05 from ANXA1sp-, I/R group).

### ANXA1sp prevents kidney tubule apoptosis/cell death

We next wanted to determine whether ANXA1sp prevented cell death in the kidney following ischemia. We performed TUNEL staining on kidney sections following severe ischemia. We found that kidneys from the ANXA1sp-treated mice had fewer TUNEL-positive nuclei following I/R, particularly at the corticomedullary junction (Fig. 2A-C). These results indicate that ANXA1sp is able to ameliorate cell death in the kidney following ischemia injury.

**Figure 2.**
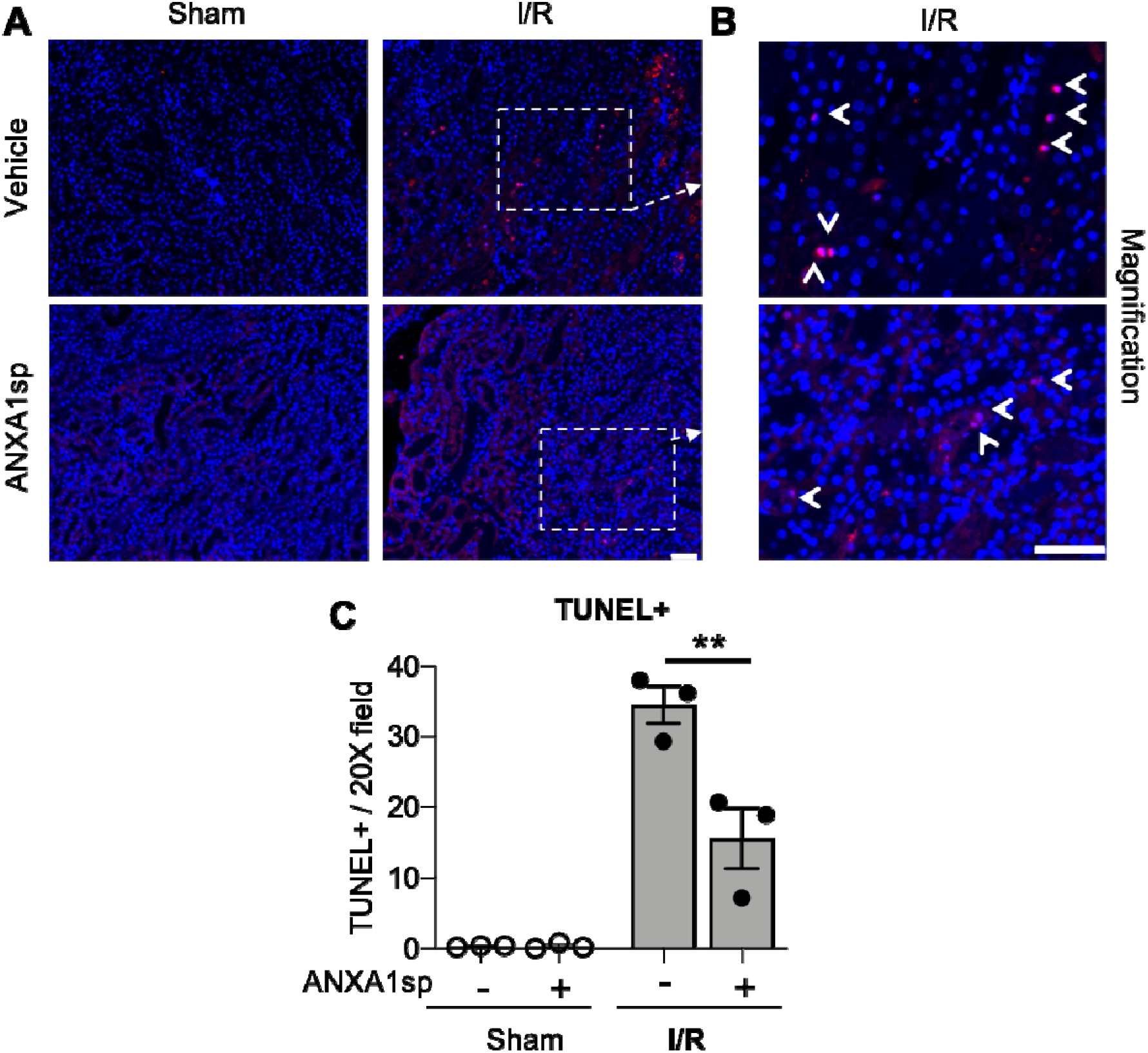
ANXA1sp treatment prevents renal cell death. Mice were treated with either Vehicle or ANXA1sp 1 hour prior to ischemia, subjected to 33 minutes of unilateral ischemia and contralateral nephrectomy and then re-injected with Vehicle or ANXA1sp 1 hour after reperfusion. (A) Kidney tissues were harvested at 24 hours after reperfusion. Representative terminal deoxynucleotidyl transferase dUTP nick end labeling (TUNEL)-stained kidney sections demonstrating increased apoptosis in Vehicle, I/R group at Day 1 after ischemia. Scale bar shows 25 um. (B) Increased magnification of boxes in (A) to demonstrate increased evidence of apoptosis in Vehicle, I/R mice (arrowhead: TUNEL-positive nuclei) compared to ANXA1sp-treated mice. Scale bar shows 25 um. (C) Quantification of TUNEL positive nuclei from (A). Graph displays mean +/− SEM of % TUNEL positive nuclei from 5 fields/section from each mouse (n=3 for Sham and I/R groups; **p<0.01).

### ANXA1sp treatment limits oxidative damage and improves mitochondrial integrity

Reactive oxygen species (ROS) are critical triggers of mitochondrial damage and can subsequently cause cell death. The 8-OHdG adduct is a product of mitochondrial (mt)DNA oxidation, and if not repaired, may lead to mutations and a dysfunctional mtDNA genome. We evaluated 8-OHdG levels in the kidney by immunofluorescence staining. Compared to sham animals, we found that 8-OHdG levels increased following ischemia (Figure 3A—compare sham to I/R panels). We found ROS staining to be localized to mitochondria as demonstrated by co-localization with citrate synthase (green) staining (Figure 3A). Following ischemia, ANXA1sp decreased the intensity of 8-OHdG staining (Figure 3A— bottom right) compared to vehicle treated mice (Figure 3A—top right), indicating that ANXA1sp limited accumulation of ROS following ischemia. In support of this notion, we found that ANXA1sp significantly increased protein levels of the antioxidant enzyme superoxide dismutase 2 (SOD2) under both sham and ischemia conditions (Figure 3B), indicating that ANXA1sp is able to upregulate protective antioxidant enzymes. To further evaluate integrity of the cellular machinery involved in maintaining the mitochondrial genome, we examined changes in protein expression of the mtDNA base-excision repair enzyme OGG1. ANXA1sp significantly increased levels of kidney OGG1 protein expression compared to vehicle treatment (Figure 3C). Moreover, as mitochondrial stress and other physiological events can promote organelle fragmentation and limit function (36), we examined the impact of ischemia on mitochondrial morphology by measuring Drp1, which is expressed in the cytosol and is recruited to mitochondria undergoing fragmentation or damage. Following ischemia, kidneys from vehicle-treated mice showed a significant increase in Drp1 protein, which was completely abrogated by ANXA1sp treatment (Figure 3D). Taken together, ANXA1sp limits ROS, promotes integrity of the mitochondrial genome, and decreases markers associated with mitochondrial fragmentation and damage.

**Figure 3.**
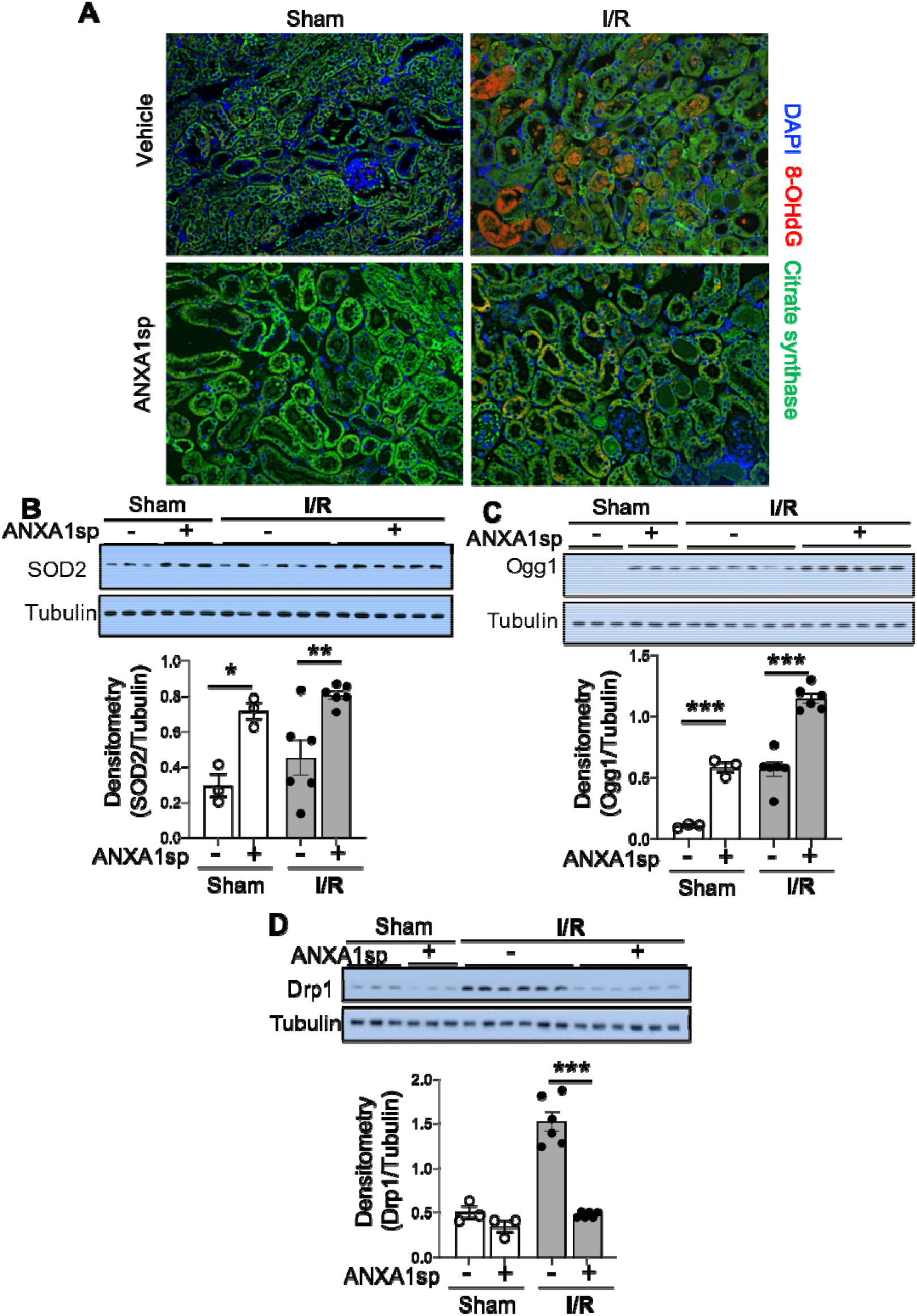
ANXA1sp treatment limits oxidative damage and improves mitochondrial integrity. Mice were treated with either Vehicle or ANXA1sp 1 hour prior to ischemia, subjected to 33 minutes of unilateral ischemia and contralateral nephrectomy and then re-injected with Vehicle or ANXA1sp 1 hour after reperfusion. Kidney tissues were harvested at 24 hours after reperfusion. (A) Representative immunofluorescence histologic staining for 8-OHdG and citrate synthase. (B) Protein levels of superoxide dismutase 2 (SOD2) were determined by Western blot with densitometry shown in graph below. (C) Protein levels of 8-Oxoguanine DNA Glycosylase (Ogg1) were determined by Western blot with densitometry shown in graph below. (D) Protein levels of dynamin-related protein (Drp)1 were determined by Western blot with densitometry shown in graph below. Graphs in B-D displays mean +/− SEM of densitometry of protein normalized to tubulin. Statistical significance determined by two-way ANOVA (n=3 samples for Sham groups, n=6 samples for I/R groups; *p<0.05, ***p<0.001).

### ANXA1sp treatment improves markers of mitophagy

The elimination of damaged mitochondria through a process termed mitophagy is also vital to maintaining cellular function in the face of cellular stress (32). Thus, we next determined the effect of ANXA1sp on mitophagy following ischemic AKI. ANXA1sp induced an accumulation of the lipidated form of microtubule associated protein 1 light chain 3 beta (LC3 II/LC3 I), a marker for autophagy (Figure 4A,B). To specifically evaluate mitophagy, we assessed levels of mitophagy regulators PINK1 (PTEN induced putative kinase 1) and Parkin. ANXA1sp treatment significantly increased both PINK1 and Parkin levels following ischemia (Figure 4C-E). We concluded that ANXA1sp improves mitophagy as an additional mechanism of kidney protection.

**Figure 4.**
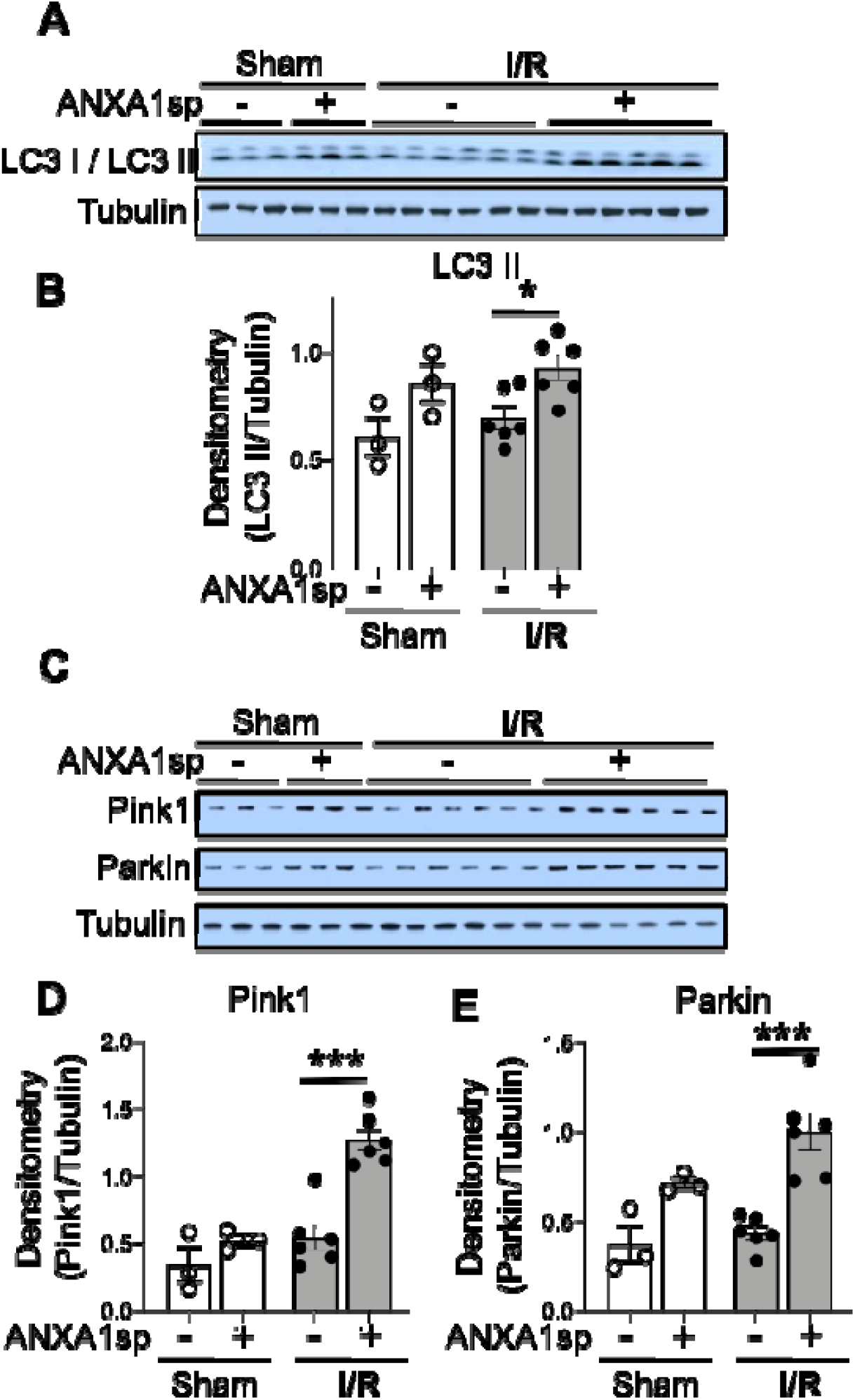
ANXA1sp treatment promotes mitophagy. Mice were treated with either Vehicle or ANXA1sp 1 hour prior to ischemia, subjected to 33 minutes of unilateral ischemia and contralateral nephrectomy and then re-injected with Vehicle or ANXA1sp 1 hour after reperfusion. Kidney tissues were harvested at 24 hours after reperfusion. (A) Western blot for LCI/III showing protein levels of indicated proteins with densitometry shown below in (B). (C) Western blot for Pink1, and Parkin with densitometry shown in (D) and (E), respectively. Graphs display mean +/−SEM of densitometry of protein normalized to tubulin. Statistical significance determined by two-way ANOVA (n=3 samples for Sham groups, n=6 samples for I/R groups; *p<0.05, ***p<0.001).

### ANXA1sp induces PGC1α-mediated mitochondrial biogenesis in the kidneys

To gain further insight into the potential impact of ANXA1sp on mitochondrial function following IR, and to determine whether ANXA1sp-mediated mitochondrial integrity is coordinated with mitochondrial biogenesis, mitochondrial biogenesis was tracked in mouse kidneys by immunoblotting for key mitochondrial proteins. Following ischemia, we found significant upregulation of the major regulator of mitochondrial biogenesis, PGC1-α (43, 44), in ANXA1sp-treated mice compared to vehicle-treated mice (Figure 5A,B). As additional evidence of improved mitochondrial biogenesis following ANXA1sp treatment, compared to vehicle, ANXA1sp treatment increased mitochondrial mass in the kidney as measured by citrate synthase (mtCS) (Figure 5C,D), cytochrome c oxidase subunit 1 of mitochondrial Complex IV (mtCI) (Figure 5C,E), and the mitochondrial NADH-ubiquinone oxidoreductase chain 1 (mtND1) (Figure 5C,F). We further found increased levels of the PGC1α target, mitochondrial transcription factor A (Tfam) (Figure 5G,H), which is required for mtDNA transcription and replication. Thus, ANXA1sp increases PGC1α, which upregulates key proteins involved in mitochondrial biogenesis, further supporting its role in mitochondrial protection following ischemic AKI.

**Figure 5.**
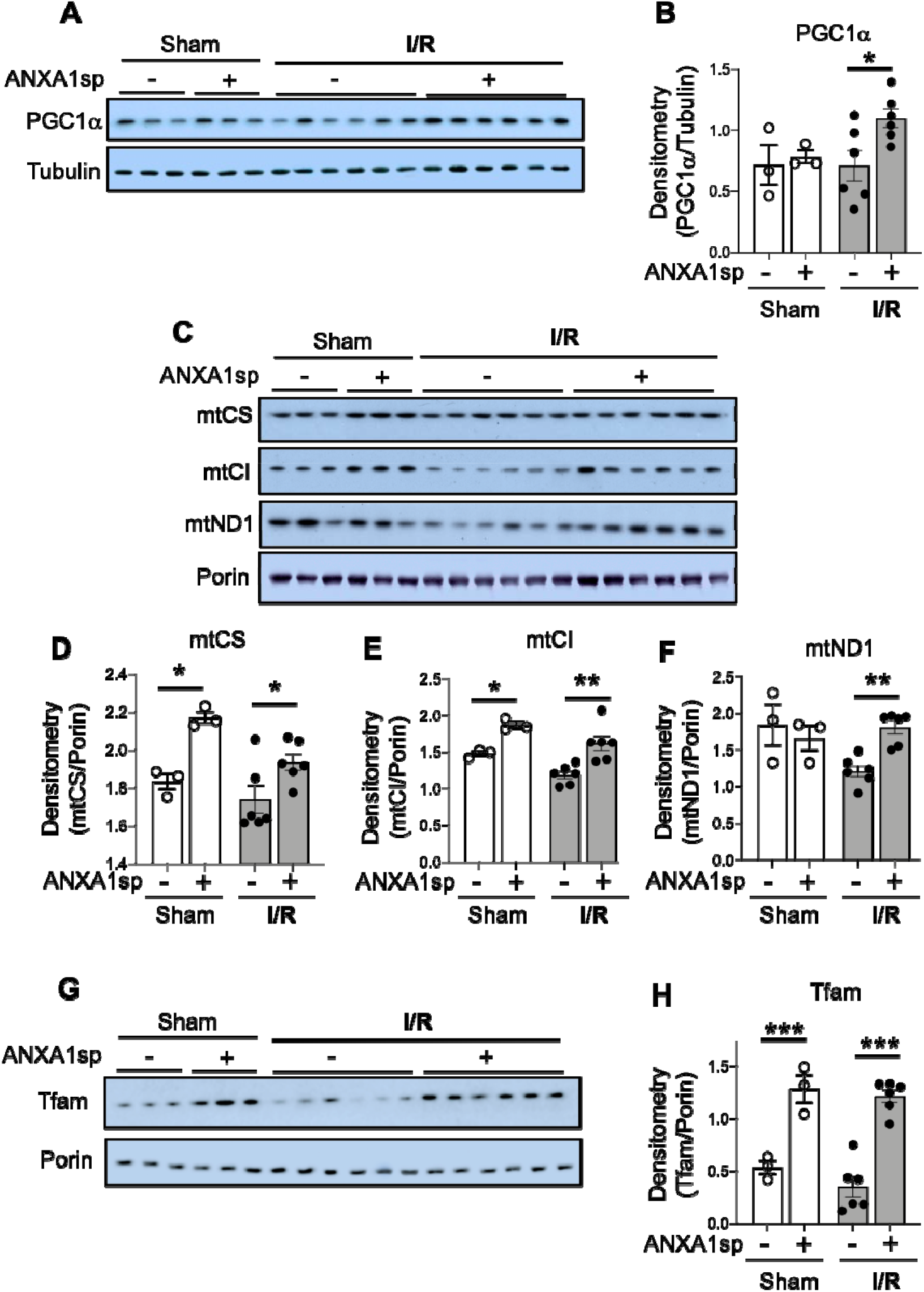
ANXA1sp treatment promotes mitochondrial integrity. Mice were treated with either Vehicle or ANXA1sp 1 hour prior to ischemia, subjected to 33 minutes of unilateral ischemia and contralateral nephrectomy and then re-injected with Vehicle or ANXA1sp 1 hour after reperfusion. Kidney tissues were harvested at 24 hours after reperfusion. (A)Western blot showing protein levels of PGC1α with densitometry shown in (B). (C) Western blot showing indicated proteins with densitometry results for (D) mitochondrial citrate synthase (mtCS), (E) mitochondrial complex I (mtCI), and (F) mitochondrial NADH-ubiquinone oxidoreductase chain 1 (mtND1). (G) Western blot showing protein levels of mitochondrial transcription factor A (Tfam) with densitometry in (H). Graphs display mean +/− SEM of densitometry of mitochondrial protein normalized to porin. Statistical significance determined by two-way ANOVA (n=3 samples for Sham groups, n=6 samples for I/R groups; *p<0.05, **p<0.01, ***p<0.001).

### ANXA1sp upregulates sirtuin-3 (SIRT3) following ischemia

We next wanted to determine cellular mediators through which ANXA1sp and PGC1α could be working to protect mitochondria and prevent cell death. SIRT3 is a mitochondrial protectant that protects against kidney injury (26, 54). SIRT3 is also a target gene of PGC1α (18). We found that ischemia caused downregulation of SIRT3 at the mRNA and protein level, an effect that was reversed by ANXA1sp treatment (Fig. 6A,B). By immunofluorescent staining, we also showed that kidneys from ANXA1sp-treated mice displayed increased SIRT3 expression after I/R compared to Vehicle, and that much of the SIRT3 expression appeared to be localized to mitochondria in kidney tubular cells as demonstrated by co-staining with mitochondrial complex IV (Fig. 6C). Thus, we concluded that ANXA1sp promotes expression of SIRT3 in mitochondria of kidney tubules following I/R, which likely helps to prevent cell death and kidney injury.

**Figure 6.**
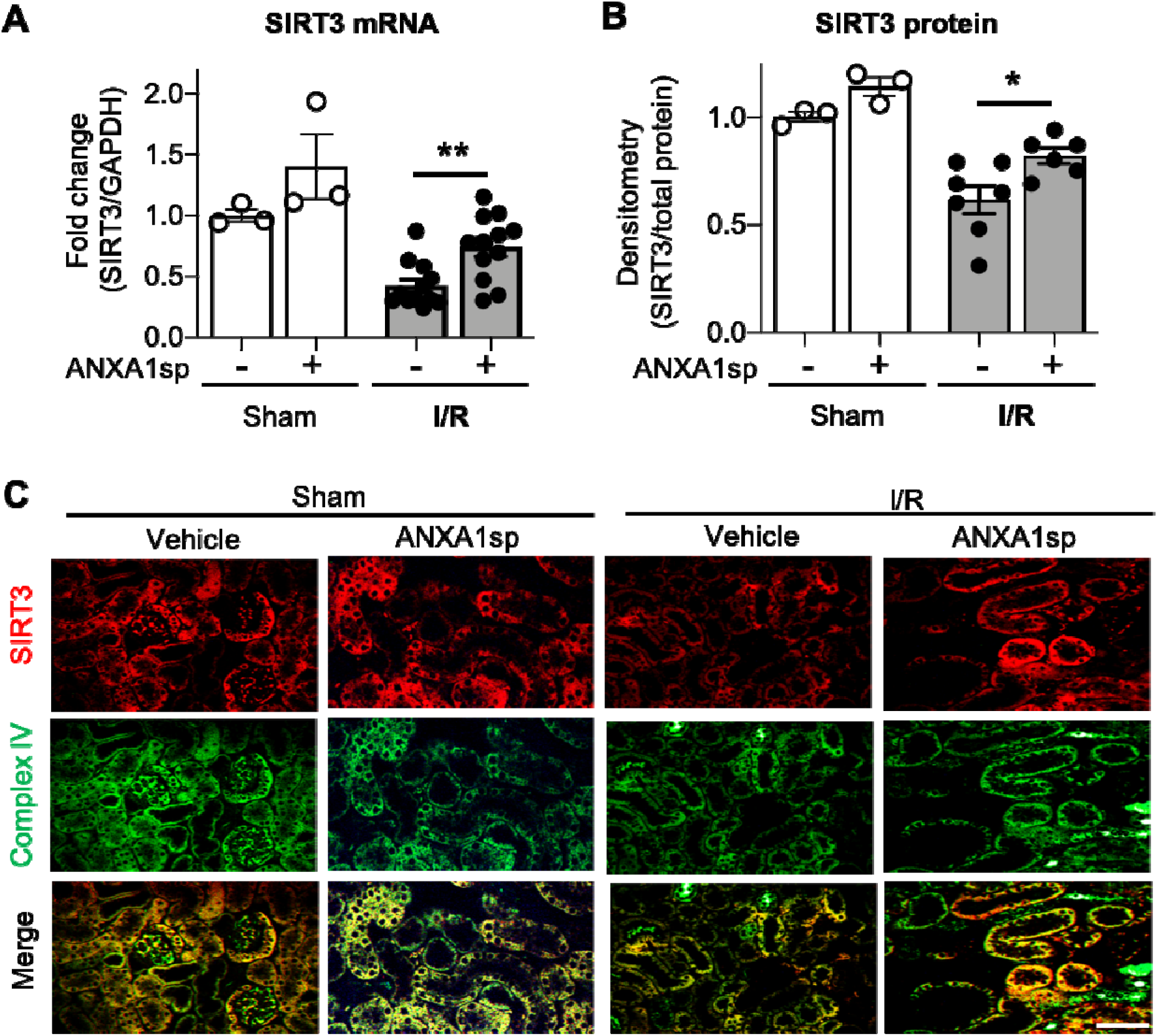
ANXA1sp treatment upregulates levels of sirtuin3 (SIRT3) in the mitochondria of kidney tubules. Mice were treated with either Vehicle or ANXA1sp 1 hour prior to ischemia, subjected to 33 minutes of unilateral ischemia and contralateral nephrectomy and then re-injected with Vehicle or ANXA1sp 1 hour after reperfusion. Kidney tissues were harvested at 24 hours after reperfusion. (A) mRNA levels of *SIRT3* mRNA were determined by RT-PCR. Graph displays mean +/−SEM of SIRT3 normalized to GADPH, then normalized to Sham, Vehicle group (n=3 sample for Sham groups, n=12 samples for I/R groups). Statistical significance determined by two-way ANOVA (**p<0.01). (B) Protein levels of SIRT3 were determined by Western blot. Graph displays mean +/−SEM of densitometry of SIRT3 normalized to total protein from stain-free gel for each sample, then normalized to Sham, Vehicle group (n=3 samples for Sham groups, n=7 samples for Vehicle, I/R group, n=6 for ANXA1sp, I/R group). Statistical significance determined by two-way ANOVA (*p<0.05). (C) Paraffin-embedded kidney tissue was analyzed for SIRT3 (top) and mitochondrial complex IV (middle) expression by immunofluorescence microscopy with merged image (bottom). SIRT3 co-localizes with mitochondrial marker complex IV in kidney tubule cells. Scale bar shows 50um.

### ANXA1sp mediates kidney protection through SIRT3

We next wanted to develop a cellular model of ischemic kidney injury in order to better test molecular mechanisms of ANXA1sp-mediated kidney protection. To mimic our in vivo model, we treated an immortalized human kidney epithelial cell line, HK-2, with vehicle or ANXA1sp and subjected cells to hypoxia using oxygen-glucose deprivation (OGD) in an anaerobic chamber. We found that ANXA1sp given prior to hypoxia exposure was able to prevent hypoxic HK-2 cell death (Fig. 7A). We then silenced SIRT3 expression with siRNA (Fig. 7B). We found that silencing of SIRT3 prevented ANXA1sp-mediated protection from hypoxic cell death (Fig. 7C). Taken together, these findings demonstrate that ANXA1sp mediates its kidney protective effects through SIRT3, which promotes mitochondrial protection and prevents cell death.

**Figure 7.**
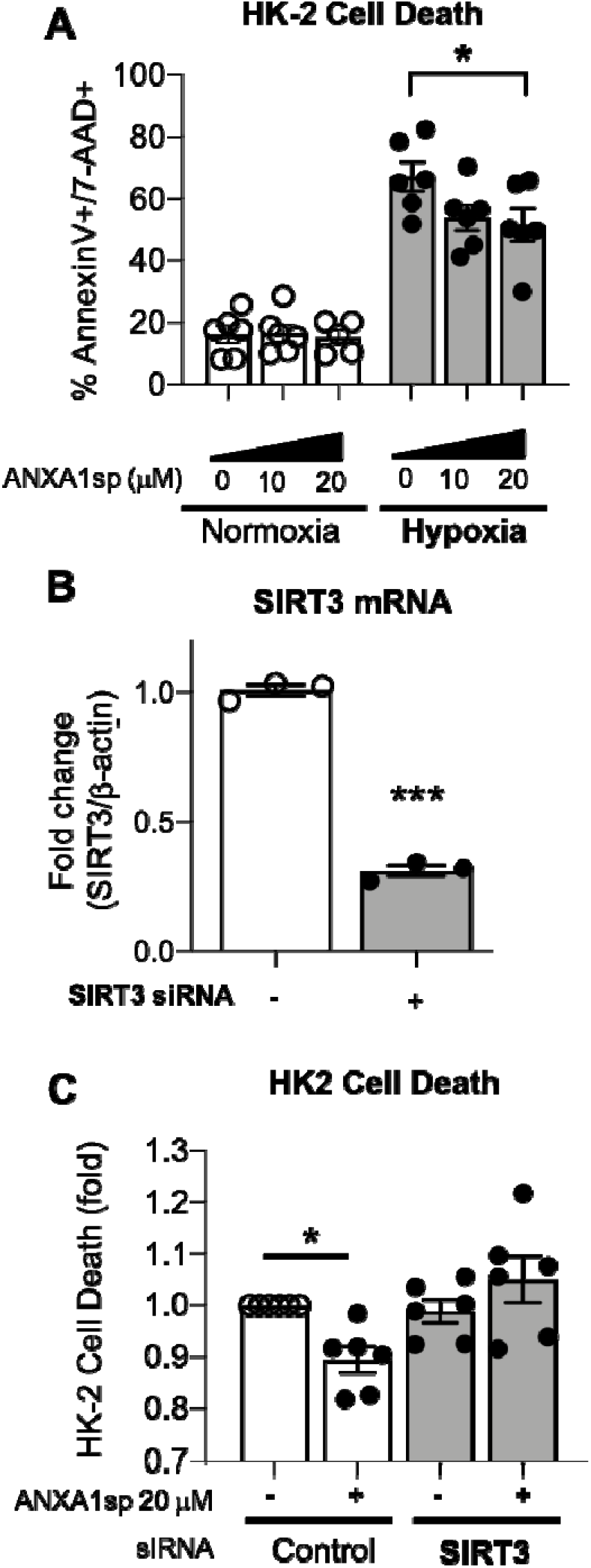
Silencing of SIRT3 prevents ANXA1sp-mediated protection from kidney cell death. (A) The immortalized human kidney cell line, HK-2, was grown to confluence in monolayers. Cells were pretreated with Vehicle or ANXA1sp at increasing concentrations and then subjected to 16 hours of oxygen-glucose deprivation (hypoxia) in an anaerobic chamber. ANXA1sp prevented hypoxic cell death (n=5-6/condition). Graphs display mean +/− SEM with significance determined by two-way ANOVA with Sidak post-hoc test (*p<0.05). (B) HK-2 cells were treated with control (-) or SIRT3 (+) siRNA. Levels of SIRT3 (A) mRNA were determined by RT-PCR. (C) HK-2 were treated with control or SIRT3 siRNA. Cells were then pre-treated with Vehicle or ANXA1sp peptide for 1hr prior to oxygen-glucose deprivation for 16 hours. Shown is fold change in cell death over Vehicle-treated, control siRNA cells. The protective effects of ANXA1sp were abrogated by SIRT3 siRNA (n=6/condition). Graph displays mean +/− SEM with significance determined by one-way ANOVA with Sidak post-hoc test (*p<0.05).

## Discussion

Our group has developed a tripeptide fragment, ANXA1sp, of the pro-resolving mediator annexin A1 that holds considerable promise for alleviating postoperative organ dysfunction. We hypothesized that ANXA1sp would alleviate AKI in a model of ischemic surgical kidney injury. In support of our hypothesis, we found that ANXA1sp limited post-surgical ischemic kidney injury. We showed that ANXA1sp treatment was associated with reduced kidney cell death following ischemia both in vitro and in vivo. We further demonstrated that ANXA1sp upregulated PGC1α and the PGC1α target SIRT3 to limit ROS, increase antioxidant enzymes, and induce mitophagy and mitochondrial biogenesis, all of which attenuated cell death. In support of this mechanism of cellular protection, we further showed that the protective effects of ANXA1sp treatment were ameliorated when SIRT3 was silenced in kidney tubule cells in vitro. Taken together, ANXA1sp augments SIRT3 levels to improve mitochondrial function, which ameliorates ischemic AKI.

AKI is a major cause of perioperative morbidity and mortality (15). Indeed, the kidney is one of the organs at particularly increased risk of injury during the perioperative period. Since the kidneys receive nearly 25% of cardiac output, they are continuously exposed to nephrotoxins during surgery (13, 49); and hemodynamic alterations, including hypotension and hypovolemia, place the kidneys at additional risk for injury. Surgeries performed on the aorta, such as abdominal aortic repair, require clamping of the abdominal aorta and/or renal vessels, which directly induces ischemic injury. Likewise, in kidney transplantation, the renal vessels are clamped to allow removal of the donor kidney. During all of these procedures, the timing of kidney insult is known. As such, the identification of kidney-protective therapeutics that can be given prior to known kidney insult holds considerable promise for limiting perioperative kidney injury. We show here that a tripeptide fragment of the parent annexin A1 molecule, ANXA1sp, limits renal damage in an ischemic kidney injury model showing its promise as a perioperative organ protectant. In addition, though it remains to be tested, since transplanted donor kidneys undergo a period of ischemia between harvest and transplant, ANXA1sp could also be used to preserve donor kidneys prior to transplantation.

The efficacy of ANXA1sp to limit organ injury in the brain (22, 53), and now the kidney, points to its mechanism of protection being modulation of a global cellular function. One possibility we considered could be limiting inflammation as the parent annexin A1 molecule is known reduce leukocyte tethering and transmigration and promote phagocytosis of dead cells (1, 7, 23, 34). However, as ANXA1sp has shown promise to limit cell death in the brain (22, 53), we tested the ability of ANXA1sp to limit cell death in the kidney following ischemia. We indeed found that ANXA1sp was able to mitigate cell death following ischemia in the kidney and in an immortalized human kidney cell line in an in vitro model of hypoxia. Taken together with its effects on limiting cell death in the brain, ANXA1sp appears to broadly limit cell death in multiple tissues.

ANXA1sp appears to limit cell death through SIRT3-mediated mitochondrial protection. We found that ANXA1sp augments expression of SIRT3 following ischemia. SIRT3 is a mitochondrial protectant that is known to promote mitochondrial biogenesis, mitophagy, metabolic flux of both carbohydrate and fatty acid substrates, and limit oxidative stress (25). ANXA1sp was able to limit cell death induced by hypoxia in vitro, and its protection was abolished when SIRT3 was silenced in HK-2 cells, implying SIRT3 is required for ANXA1sp-mediated kidney protection. Our data showing augmented SIRT3 expression as a kidney protectant is in line with other groups that have shown that SIRT3 deletion is deleterious in both nephrotoxic and septic AKI (26, 54). Furthermore, ANXA1sp also benefits mitochondrial function in the ischemic kidney by promoting mitochondrial biogenesis. Our data show that PGC-1α, a major transcriptional co-activator for mitochondrial biogenesis (45), is upregulated by ANXA1sp following ischemia. PGC-1α is known to limit AKI (35, 43, 44). PGC1-α is also required for the induction of many ROS-detoxifying enzymes, including SOD2 and catalase (37), and is also known to induce SIRT3 expression (18).

Owing to its mitochondrial protective properties, ANXA1sp appears to be particularly efficacious at limiting cell stress in tissues with high mitochondrial content. Due to the high metabolic demands required to remove waste products and regulate acid-base status, fluids, electrolytes, and blood pressure, the kidney is one of the most metabolically active organs in the body. In fact, of all organs, the kidney has the second highest mitochondrial content and oxygen consumption after the heart (27, 28). Damaged mitochondria increase ROS production, propagating mitochondrial damage (11), which may stimulate the release of pro-cell death factors, such as cytochrome c, from mitochondria into cytosol to initiate cell death (41). Because ischemia-reperfusion events are frequently complicated by mitochondrial dysfunction and increased ROS production, we tested the ability of ANXA1sp to limit these deleterious events. We showed that ANXA1sp limits cell death and ROS production and increases antioxidant enzymes. ANXA1sp suppression of ROS in the kidney is again likely due to its upregulation of SIRT3 (31): We show that ANXA1sp upregulates OGG1, an important mitochondrial DNA repair enzyme, and SIRT3 can interact with OGG1 (6). In addition, under stress conditions, ROS production can exceed the antioxidant capacity of the mitochondria due to acetylation-induced inactivation of mitochondrial antioxidant enzymes SOD2, which is deacetylated and activated by SIRT3 (9). Thus, in addition to the direct upregulation of SOD2 expression that we show here, ANXA1sp-mediated upregulation of SIRT3 could also augment activation of SOD2 to limit ROS damage. Taken together, ANXA1sp augments the mitochondrial antioxidant system to prevent ROS-induced mtDNA oxidation and damage.

ANXA1sp also improves mitophagy, a selective form of autophagy that eliminates redundant or damaged mitochondria (14). Recent evidence suggests that mitophagy plays an important role in AKI development and subsequent kidney repair: PINK1-Parkin-mediated mitophagy was reported to be protective against both toxic and ischemic kidney injury (40, 46). Not only did ANXA1sp treatment improve mitophagy in our studies, but it also reduced mitochondrial-associated Drp1, suggesting that ANXA1sp limits mitochondrial fragmentation. The influence of Drp1 and mitochondrial fragmentation on apoptosis and exacerbation of injury has been documented in several studies (3-5), and inhibiting this response likely protects the kidney from further injury after ischemia. The specific mechanism by which ANXA1sp decreases mitochondrial-associated Drp1 is not known; however, activation of the SIRT3 and mitochondrial biogenesis and mitophagy program may also influence mitochondrial dynamics after ischemia.

The mechanisms by which ANXA1sp upregulates SIRT3 to protect the mitochondria are still unclear. The parent annexin A1 molecule binds the formyl peptide receptor 2 (FPR2), a promiscuous G-protein coupled receptor (GPCR) that serves as a pattern recognition receptor for bacterial formylpeptides, eicosanoid lipid molecules and pro-resolving lipid mediators (55). As ANXA1sp is a small tripeptide fragment of the parent annexin A1 molecule, it is likely that ANXA1sp is able to bind to and activate FPR2. However, due to its small size, ANXA1sp may have multiple cellular targets. The identification of the cellular target of ANXA1sp is an active area of investigation in our laboratory.

In conclusion, the annexin A1 mimetic peptide, ANXA1sp, upregulates SIRT3 and improves mitochondrial function in the kidney following ischemic injury. Restoration of mitochondrial function via ANXA1sp/SIRT3 may offer unique pharmacological targets for improved recovery from AKI. Thus, ANXA1sp holds considerable promise as a perioperative kidney protectant.

## Author contributions

H.S. helped with study design, conducting experiments, acquiring data, providing reagents, writing/editing manuscript. Q.M. helped with study design, conducting experiments, acquiring data, editing manuscript. Z.Z. helped with study design, conducting experiments, acquiring data, providing reagents, editing manuscript. J.R helped with acquiring data, editing manuscript. B.T.M. helped with conducting experiments, acquiring data, editing manuscript. S.D.C. helped with study design, providing reagents, editing manuscript. L.U. helped with study design, editing manuscript. J.R.P helped with study design, conducting experiments, acquiring data, providing reagents, writing manuscript.

## Acknowledgements

We thank Dr. Paul Wischmeyer for allowing use of anaerobic chamber for in vitro experiments. Funding: This work was supported by National Institutes of Health grants K08 GM132689 to JRP and R01 DK118019 to SDC, and US Veterans Health Administration, Office of Research and Development, Biomedical Laboratory Research and Development grant BX000893 to SDC. The funding sources played no role in study design; in the collection, analysis and interpretation of data; in the writing of the report; or in the decision to submit the article for publication.

